# Individuals with Intermittent Explosive Disorder Exhibit Idiosyncratic Neural Responses during Social-emotional Processing

**DOI:** 10.64898/2026.03.13.711681

**Authors:** Jiajie Chen, Sarah Keedy, Emil Coccaro, Yuan Chang Leong

## Abstract

Intermittent explosive disorder (IED) is associated with impulsive aggression in ambiguous social contexts. Prior neuroimaging studies have treated IED as a homogenous group, but identical social situations may elicit divergent responses across IED individuals. Here, we test the hypothesis that IED is characterized by idiosyncratic neural responses to social cues during naturalistic social-emotional processing. IED individuals and healthy controls completed a validated paradigm where they were presented with video vignettes of interpersonal interactions while undergoing fMRI. We computed the intersubject correlation (ISC) in neural time courses between pairs of participants to quantify neural similarity, and assessed whether similarity differed between Healthy-Healthy and IED-IED dyads using Bayesian multilevel models, controlling for self-reported emotional responses and intention attributions for each vignette. Healthy-Healthy dyads showed significantly higher ISC than IED-IED dyads, indicating that neural responses to the videos were similar among healthy participants, but idiosyncratic in IED individuals. These effects were observed in regions in the default mode and salience networks, including the precuneus, medial prefrontal cortex, superior temporal sulcus, insula, and dorsal anterior cingulate cortex. Individuals with IED exhibited idiosyncratic neural responses during naturalistic social-emotional processing, even after accounting for differences in emotional reaction and intention attribution. This neural idiosyncrasy may reflect atypical integration of social cues, giving rise to maladaptive interpretations and impulsive aggression. Assessing neural synchrony during ecologically valid paradigms offers a promising tool for identifying neural markers of interpersonal dysfunction and informing targeted interventions.

---

Aggression, defined as social behaviors intended to harm another individual (Baron, 1977; Baron & Richardson, 1994; Bushman & Huesmann, 2010), emerges from social-emotional processes that transform cues about other people’s intentions into affect, appraisal, and action (Crick & Dodge, 1994, 1996). These processes are disrupted in intermittent explosive disorder (IED), a condition characterized by recurrent episodes of impulsive aggression (Coccaro, 2011, 2012; Coccaro, Fanning, et al., 2016). Affecting approximately 4.0% of the population, IED is associated with heightened risk of intimate partner violence, gun use, and other aggressive behaviors (Coccaro & Lee, 2020). These maladaptive behaviors often emerge in response to perceived social threats, underscoring the importance of understanding how individuals with IED interpret social situations. Social-emotional information processing (SEIP) refers to the set of cognitive and affective processes by which individuals perceive, interpret, and respond to others’ behavior (Lemerise & Arsenio, 2000). Within this framework, individuals with IED report stronger negative emotional reactions to ambiguous situations and are more likely to attribute others’ actions to deliberate harm (Coccaro et al., 2009; Coccaro, Fanning, & Lee, 2017; McCloskey et al., 2006). This perceived hostility can in turn trigger impulsive reactions, increasing the likelihood of reactive aggression.

IED and aggressive behavior have been linked to alterations across multiple neural systems and brain regions (Blair et al., 2025; Paliakkara et al., 2025). Prior neuroimaging studies point to consistent disruptions in the brain circuits that regulate emotion and social evaluation in IED. Specifically, individuals with IED show reduced gray matter volume in orbitofrontal and ventromedial prefrontal cortex, anterior cingulate cortex, amygdala, and insula compared with both healthy and psychiatric controls (Coccaro, Fitzgerald, et al., 2016; Seok & Cheong, 2020). IED individuals have also been shown to exhibit heightened amygdala reactivity and reduced orbitofrontal cortex activity when viewing angry facial expressions compared with healthy controls (Coccaro et al., 2007). More recently, Coccaro and colleagues (Coccaro et al., 2022) found that individuals with IED showed reduced activation in medial orbitofrontal and anterior cingulate cortices when viewing interactions that could be interpreted as aggressive compared with clearly benign interactions, relative to healthy controls.

The past studies, however, have focused on group-level differences, comparing IED individuals with healthy controls. Implicit in this approach is the assumption that individuals with IED form a homogeneous group, responding in broadly similar ways to the same social stimuli. Yet growing evidence suggests that negative psychological traits are often marked by idiosyncrasy. For example, social stimuli elicit idiosyncratic neural responses in lonely individuals, whereas non-lonely individuals were more aligned (Baek et al., 2023; Broom et al., 2024). Similarly, pessimists exhibit variable neural responses when imagining future events, whereas neural responses of optimists were similar to one another (Yanagisawa et al., 2025). These findings suggest that negative dispositions may be characterized not only by deviations from healthy controls, but also by the uniqueness of each individual. This pattern has been termed the Anna Karenina effect, echoing the opening line of Tolstoy’s novel: “Happy families are all alike; every unhappy family is unhappy in its own way” (Berman et al., 2013; Diamond, 1997; Finn et al., 2020).

Consistent with the Anna Karenina effect, individuals with IED may each develop idiosyncratic “scripts” shaped by their personal histories, such that different cues trigger threat perceptions in different people. In this view, IED individuals would exhibit divergent, individualized neural responses to the same social situation, in contrast to healthy individuals who tend to rely on shared social schemas that promote convergent interpretations. The opposite pattern, however, is also plausible: individuals with IED may converge on a common hostile interpretation of ambiguous cues, whereas healthy individuals may vary in how they explain away potentially threatening situations (Finn et al., 2018; Iyer et al., 2023).

To test these possibilities, we examined neural synchrony during naturalistic social-emotional processing in individuals with IED and healthy controls. Participants completed a validated video-based SEIP paradigm (Coccaro et al., 2021, 2022; Coccaro, Fanning, Fisher, et al., 2017), in which they viewed videos of everyday interpersonal interactions varying in ambiguity, while undergoing fMRI. We computed the intersubject correlation (ISC) of neural time courses to quantify the extent to which participants’ brain responses converged during video viewing (Lyu et al., 2024; Nastase et al., 2019). We hypothesized that healthy controls would exhibit greater neural synchrony with one another, whereas individuals with IED would show more idiosyncratic neural responses, consistent with an Anna Karenina effect. Alternatively, if individuals IED converge on a consistent interpretative bias, we would expect IED participants to exhibit greater synchrony with one another than with healthy controls.

## Methods

### Participants

45 right-handed adult participants were recruited through advertisements and flyers targeting both those with problematic anger or aggression and healthy controls. Participants with a life history of bipolar disorder, schizophrenia (or other psychotic disorder), or intellectual disability were excluded from this study, as were those with a current history of alcohol or other substance use disorders. All participants were medically healthy and medication free, and provided informed consent prior to the experiment. Study procedures were approved by the IRB at the University of Chicago. These data have been previously published in (Coccaro et al., 2022), however, all analyses reported in this study are new.

### Diagnostic and Trait Assessments

Diagnoses were determined according to DSM-5 criteria (American Psychiatric Association, 2013) by individuals with graduate degrees in mental health. 19 participants were diagnosed with IED, while the remaining 26 participants were designated as healthy controls. Additional diagnoses for IED participants are displayed in Table S1. All participants were assessed for aggression using the Life History of Aggression (LHA) measure (Coccaro et al., 1997).

### The Video Social-Emotional Information Processing (V-SEIP) Task

The Video Social-Emotional Information Processing (V-SEIP) task consists of 40 trials. Each trial included three phases: Video, Attribution, and Emotional Response, separated by a jittered fixation period of 4-12 seconds (Figure 1).

**Figure 1.**
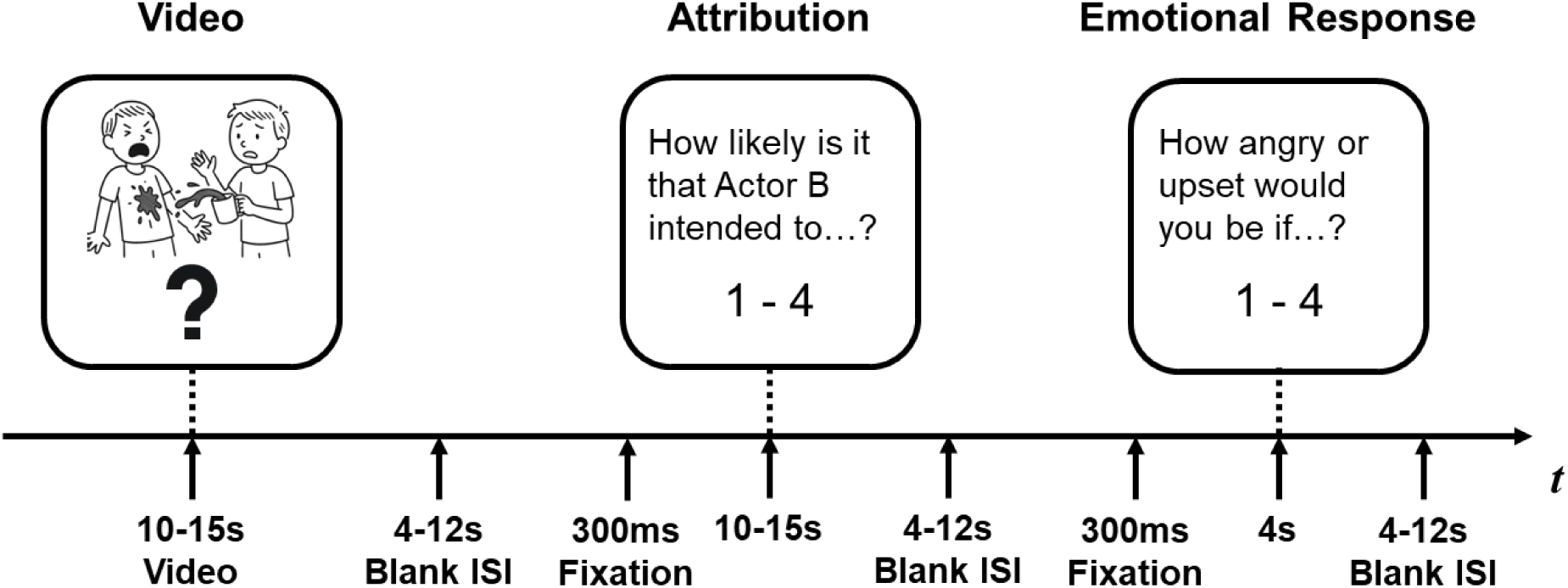
The V-SEIP Task. The task consists of three consecutive phases: the Video phase, where participants watch a video depicting a social situation; the Attribution phase, where participants report their attribution of the actor’s intent; the Emotional response phase, where participants report their emotional response.

During the Video phase, participants were presented with one video (duration: 10-15 seconds) depicting a social interaction between two individuals. There were 10 unique situations depicted in the videos, each with two possible endings: socially adverse but ambiguous in intent (adverse videos) or clearly non-aggressive (control videos). Adverse and control videos were identical until the final 2 seconds. For example, in one scenario, a hotel guest either accidentally spills coffee on another person’s shirt (adverse ending) or simply pours their own coffee without incident (control ending). Half of the scenarios were related to physical actions (e.g., spilling coffee), and the other half were related to relational actions (e.g., betraying a coworker’s confidence). All situations were filmed with both male and female actors as the primary subject, leading to a total of 40 videos (10 situations × 2 endings × 2 sexes of primary subject).

Following the Video phase, the Attribution phase assessed participants’ interpretation of the primary subject’s intent. A written question was displayed on the screen for 4 seconds: “How likely is it that Actor B intended to hurt or embarrass Actor A in the video?” Participants responded using a 4-point scale (1 = “not at all” to 4 = “very”). This was followed by the Emotional Response phase, which measured participants’ emotional reactions to the event. A second written question was displayed for 4 seconds: “How angry or upset would you be if this happened to you?” Participants again rated their response on the same 4-point scale. In the study, the order of the Attribution phase and the Emotional Response phase was pseudo-randomized.

The 40 trials were presented over four runs in a single fMRI scan session. Each run in the scanning session consists of five trials with adverse video and five trials with control video. Consistent with past work using the V-SEIP (Coccaro et al., 2009), the trial order was preserved for all participants.

### fMRI Data Acquisition and Preprocessing

Imaging was performed using a Philips Achieva Quasar 3T MRI Scanner. To facilitate anatomical landmark identification and alignment with functional images, a high-resolution T1-weighted structural scan was obtained with the following parameters: repetition time (TR) of 8.0 ms, echo time (TE) of 3.5 ms, flip angle of 8°, field of view (FOV) of 240 mm, and a slice thickness of 1.0 mm with no inter-slice gap. Functional MRI data were acquired through dynamic T2*-weighted gradient echo planar imaging (EPI) sequences designed to capture BOLD (blood oxygenation level-dependent) signals. These scans used a TE of 25 ms, TR of 2000 ms, flip angle of 77°, FOV of 192 mm, and consisted of 30 oblique axial slices (4 mm thickness with a 0.5 mm gap) oriented approximately parallel to the anterior commissure-posterior commissure (AC-PC) line.

All images were preprocessed with fMRIPrep version 24.0.1 (Esteban et al., 2019). Additional high-pass filtering (0.01 Hz) was applied by clean_img function in Nilearn (version 0.10.4; Nilearn contributors et al., 2025). Detailed descriptions of the fMRIPrep preprocessing pipeline are provided in Supplemental Methods. To account for the hemodynamic delay, the stimuli were shifted by 4.5 seconds when aligning with the BOLD response. We parcellated the whole-brain neural responses by applying the 200-region cortical parcellation from Schaefer et al. (Schaefer et al., 2018), together with 21 subcortical regions defined by the Harvard-Oxford subcortical atlas (Desikan et al., 2006), resulting in a total of 221 regions of interest (ROIs). Within each ROI, voxel-wise BOLD time courses were averaged to derive the time course of the ROI.

### Neural Similarity

We computed pairwise ISC (Figure 2) at neural level during the video phase. To compute pairwise ISC, we first extracted preprocessed fMRI time series corresponding to the video phase. Within each trial, the time series was z-scored separately. We then concatenated the z-scored data across all 40 video phases for each participant within each ROI. Pearson correlations were then computed between the time series of each dyad within each ROI, yielding dyadic ISC values at the ROI level. These correlations capture the extent to which two participants show synchronized neural responses to the video stimuli within the same brain region. All ISC values were Fisher z-transformed prior to analysis.

**Figure 2.**
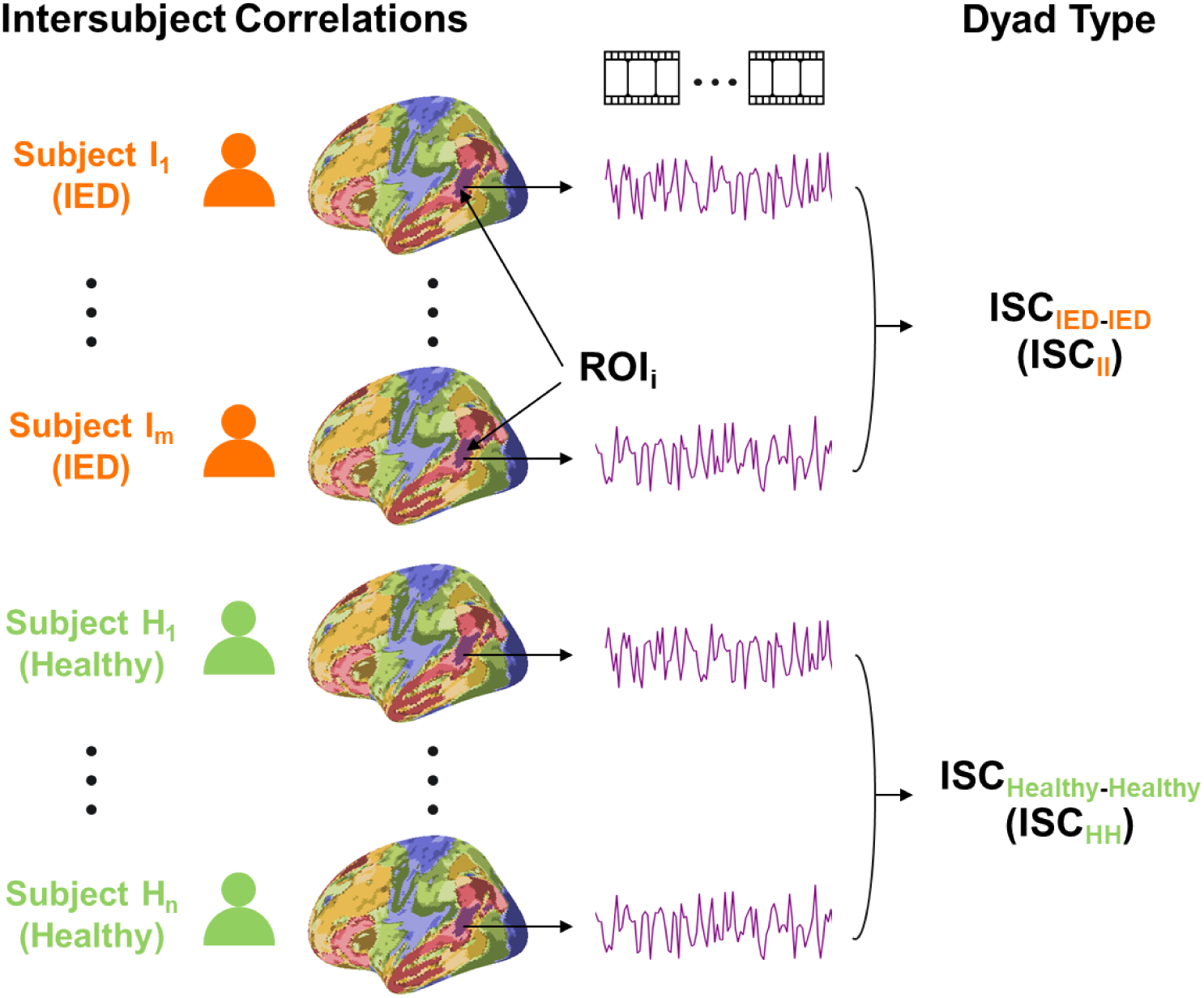
Intersubject correlation (ISC) analyses. We calculated the Pearson correlations of the neural time courses during the video phase for each dyad at each ROI. Our main analysis tests if ISC values between healthy participants (ISC_HH_) are higher than those between IED participants (ISC_II_).

### Similarity of Attribution and Emotional Response Ratings

We quantified how similarly each pair of participants explicitly evaluated the vignettes. These dyad-level measures were included as covariates to ensure that group differences in ISC were not simply attributable to idiosyncratic intentionality attributions or emotional reactions. Similarity was calculated separately for participants’ ratings in the Attribution and Emotional Response phases. For each phase, each participant’s ratings across all 40 videos were vectorized. For each dyad, we computed the Euclidean distance *d* between the rating vectors and derived behavioral similarity as 1 / (1 + *d*) such that higher values indicate more similar ratings. Missing responses were excluded pairwise so that both members of each dyad contributed ratings from the same set of videos when computing distance.

### Bayesian Multilevel Models

To test whether participants with IED exhibit more idiosyncratic neural responses, we examine group-level differences in dyadic ISCs using Bayesian multilevel models. Dyads were categorized based on the number of IED participants they contained, resulting in three distinct groups: Healthy-Healthy (HH), IED-Healthy (IH), and IED-IED (II). If IED participants are more idiosyncratic, we expect ISC values in HI and II dyads to be systematically lower than those in HH dyads.

For each ROI, we fit a Bayesian multilevel model to model ISC as a function of dyad category (HH, IH, or II). Dyadic similarity in attribution and emotion response ratings were included as covariates. A multi-membership random intercept for the two dyad members was also included to account for individual contributions to dyadic outcomes. We chose this approach because dyadic ISC values are hierarchically organized: each ISC observation is defined by a pair of participants, and multiple dyads share the same individuals. This approach accounts for the resulting non-independence among observations and directly quantifies uncertainty in all parameters via their posterior distributions (Chen et al., 2020). Group differences were evaluated through posterior contrasts between dyad types, derived directly from the estimated posterior means of the dyad type parameter. We performed three planned contrasts (1) HH vs. II, (2) HI vs. II, and (3) HH vs. HI. These posterior contrasts quantify how ISC varies as dyads include more IED participants. To assess the robustness of our results, we also repeated the analyses separately for adverse and control videos, and while controlling for the racial composition of the dyads.

We considered a parameter to show a credible effect when more than 95% of its posterior distribution lay above or below zero. To address multiple comparisons, we applied a threshold based on the expected posterior error probability (PEP), retaining effects with PEP < 0.01 (Käll et al., 2008; Storey, 2003). This corresponds to less than a 1% chance of incorrectly inferring the direction of an effect and is conceptually similar to controlling the false discovery rate (FDR) at q < 0.01 in frequentist terms. Among the ROIs with 95% highest density intervals (HDIs) above zero, we report those that remained significant after applying the PEP threshold.

Models were estimated in R (v.4.4.1) (R Core Team, 2024) using the brms package (v.2.23.0) (Bürkner, 2017). We used weakly informative priors consistent with Chen et al. (Chen et al., 2020) (see the Supplemental Methods for the prior specification). All models were estimated using four Markov chain Monte Carlo (MCMC) chains, each with 4,000 iterations including 2,000 warm-up iterations. Posterior estimates were based on the remaining 8,000 samples combined across chains. Model convergence was assessed using the Gelman-Rubin statistic (R^ < 1.1) and effective sample size (ESS > 200) for all parameters. To evaluate model fit, we compared the full model to a corresponding null model that excluded dyad type. Model performance was evaluated using approximate leave-one-out cross-validation (LOO), focusing on differences in expected log predictive density (ELPD). ROIs in which the null model showed clearly better predictive performance than the full model (|ΔELPD| > 4) (Sivula et al., 2025) were excluded from further interpretation.

## Results

### Participant Characteristics

IED participants (*M* = 17.33 ± 4.81) had significantly higher LHA aggression scores than the healthy participants (*M* = 4.24 ± 3.14; *t*(41) = −10.80, *p* < .001; Figure 3), indicating that individuals with IED reported substantially greater lifetime frequency of overt aggressive behavior. There were no significant differences in distribution in sex, age and socioeconomic status between healthy and IED groups, but the groups differed in race composition, with fewer white IED participants (Table 1).

**Figure 3.**
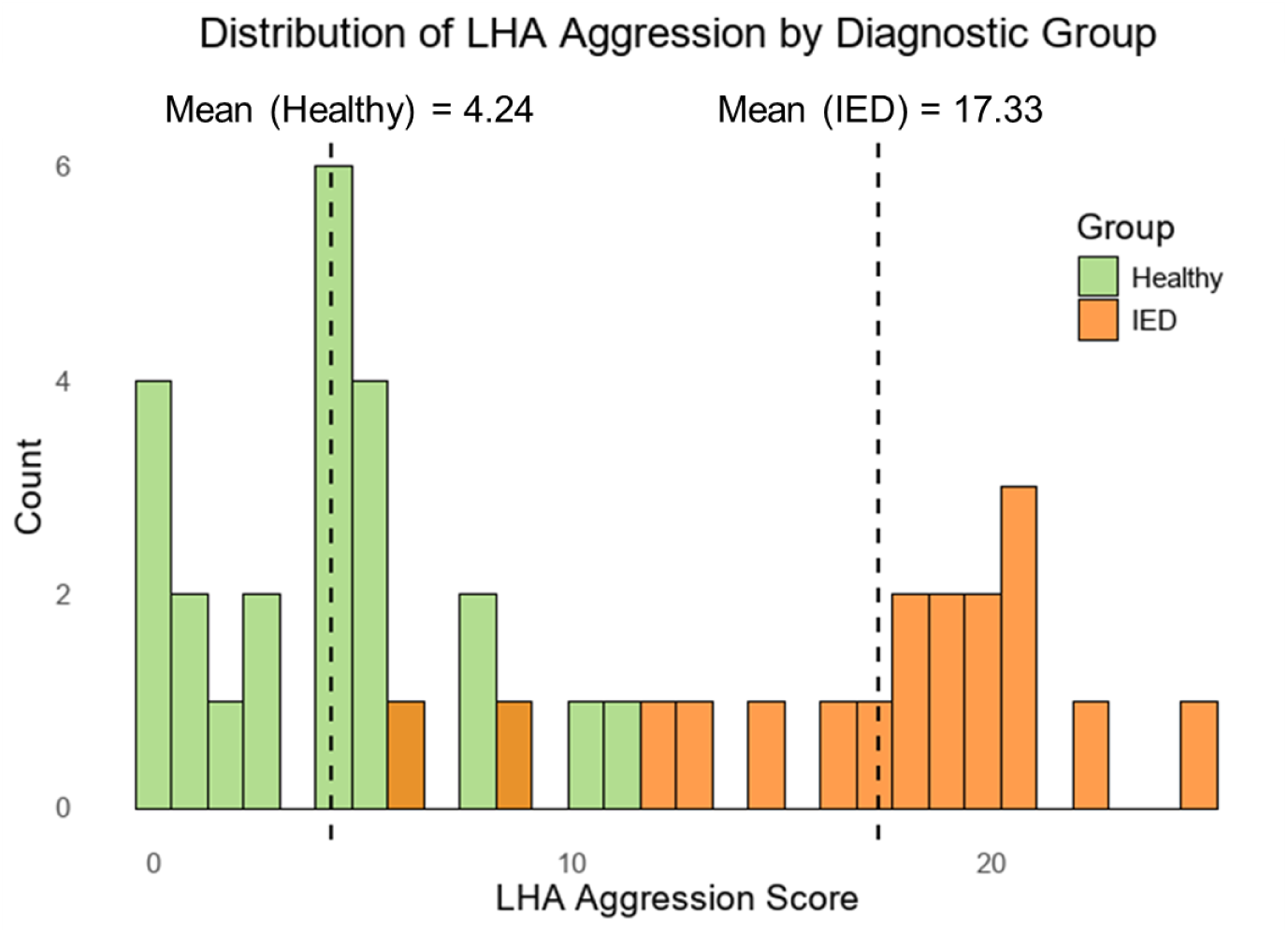
Distribution of Life History of Aggression (LHA) scores in individuals with IED patients (green) and healthy controls (red). Dotted lines indicate group mean.

**Table 1.** Participant Demographics (N = 45). Values are reported as N (percentage %) or Mean ± SD. (AA: African American; HS: high school).

|  | IED | Healthy Control | Test Statistic | *p* Value |
| --- | --- | --- | --- | --- |
| Sex, Female | 11 (58%) | 13 (50%) | $\chi^2 = 0.05$ | .824 |
| Age, Years | 35.0 ± 10.8 | 32.0 ± 8.5 | $t = -1.01$ | .318 |
| LHA Aggression | 17.33 ± 4.81 | 4.24 ± 3.14 | $t = -10.80$ | < .001 |
| SES score | 35.79 ± 14.94 | 41.54 ± 12.07 | $t = 1.43$ | .161 |
| Race (N:<br>White/AA/Other) | 2/15/2 | 12/10/4 | $\chi^2 = 7.91$ | .019 |
| Education (N:<br>HS/College/Post-C<br>ollege) | 2/16/1 | 1/22/3 | $\chi^2 = 1.22$ | .543 |

### Neural Responses to Social Situations Are Idiosyncratic in Individuals with IED

If IED patients responded idiosyncratically to the videos, dyads of two IED participants (II) would exhibit reduced ISCs compared to dyads of healthy participants (HH) and mixed dyads with one IED participant and one healthy participant (HI). To test this possibility, we fit separate Bayesian multilevel models for each of the 221 ROIs to predict dyadic ISC from dyad type.

ISC between healthy participants were higher than between IED participants (Figure 4). Specifically, ISC_HH_ was higher than ISC_II_ in the precuneus (*β* = 0.088; 95% HDI, [0.035, 0.14]; *P*(*β* > 0) = 1), insula (*β* = 0.023; 95% HDI, [0.004, 0.044]; *P*(*β* > 0) = 0.99), superior temporal sulcus (STS) (*β* = 0.18; 95% HDI, [0.041, 0.32]; *P*(*β* > 0) = 0.99), intraparietal sulcus (IPS) (*β* = 0.031; 95% HDI, [0.006, 0.057]; *P*(*β* > 0) = 0.99), dorsal anterior cingulate cortex (dACC) (*β* = 0.022; 95% HDI, [0.002, 0.042]; *P*(*β* > 0) = 0.98), dorsolateral prefrontal cortex (dlPFC) (*β* = 0.022; 95% HDI, [0.001, 0.043]; *P*(*β* > 0) = 0.98), medial prefrontal cortex (mPFC) (*β* = 0.030; 95% HDI, [0.004, 0.056]; *P*(*β* > 0) = 0.99) and nucleus accumbens (*β* = 0.028; 95% HDI, [0.008, 0.048]; *P*(*β* > 0) = 0.99).

**Figure 4.**
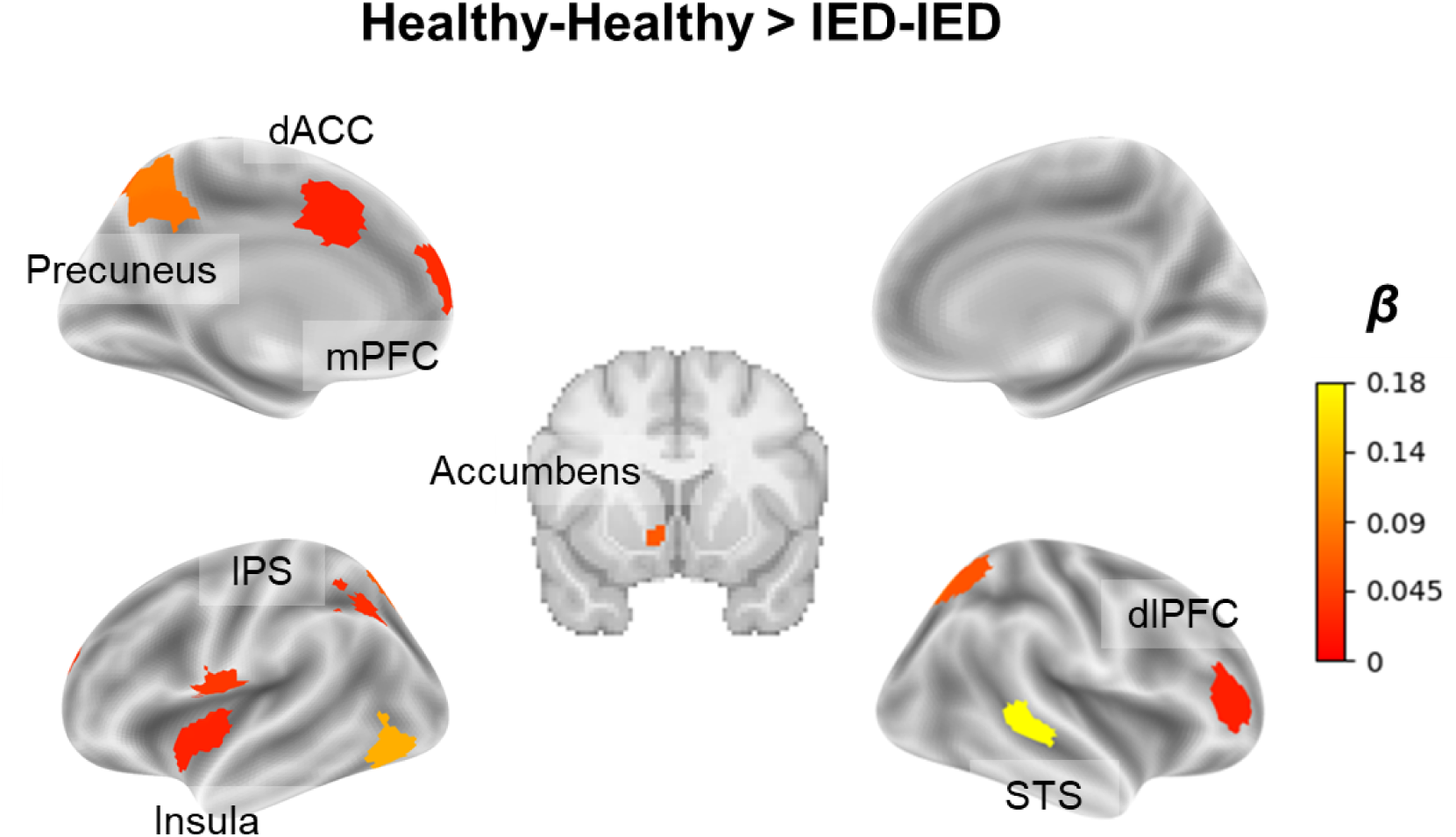
ROIs where ISC between healthy participants was higher than ISC between IED participants. Color bar indicates the estimated posterior means of the contrast. ISC_HH_ was higher than ISC_II_ in the precuneus, insula, superior temporal sulcus (STS), intraparietal sulcus (IPS), dorsal anterior cingulate cortex (dACC), dorsolateral prefrontal cortex (dlPFC), medial prefrontal cortex (mPFC) and accumbens. The model controlled for dyadic similarity in attribution and emotional responses.

A similar pattern was observed in the secondary contrasts, with ISC_HH_ higher than IS C_HI_ and ISC_HI_ higher than ISC_II_ (Supplementary Figure S1). In all cases, there were no ROIs with a credible effect in the reverse direction. Similar results were observed when repeating the analyses separately for adverse and control videos (Supplementary Figure S2). As race was confounded with IED diagnosis, we re-ran the analysis while controlling for race as a covariate. ISC remained higher between healthy participants than IED participants in the precuneus (*β* = 0.092; 95% HDI, [0.032, 0.152]; *P*(*β* > 0) = 1.00), mPFC (*β* = 0.034; 95% HDI, [0.003, 0.064]; *P*(*β* > 0) = 0.98), and insula (*β* = 0.030; 95% HDI, [0.007, 0.053]; *P*(*β* > 0) = 0.99) when controlling for the racial composition of the dyad (Supplementary Figure S3).

Together, the results provide converging evidence that individuals with IED exhibit more idiosyncratic neural responses during naturalistic social-emotional processing, such that neural time courses are less synchronized across IED participants than across healthy controls.

## Discussion

In this study, we investigated whether individuals with IED exhibit idiosyncratic neural responses to social situations. Using ISC analyses, we found that neural responses while viewing videos of social interactions were less synchronized between IED participants than between healthy participants. These effects were observed across multiple brain regions implicated in social cognition, executive control, and affective valuation, suggesting that idiosyncratic neural responses in these networks may underlie the tendency toward reactive aggression in IED. These group differences in neural synchrony survived controlling for dyadic similarity in intention attribution and emotional response ratings, suggesting that the observed neural idiosyncrasy in IED was not merely due to differences in explicit attributions of hostility or self reported emotional reactions to the videos.

The neural idiosyncrasy observed in individuals with IED was particularly evident in hubs of the default mode network (DMN), including the precuneus, superior temporal sulcus, and medial prefrontal cortex. These regions have been consistently implicated in social cognition, self referential processing, and theory of mind (Menon, 2023; Raichle, 2015; Smallwood et al., 2021; Spreng et al., 2009). Prior work has shown that DMN regions exhibit tightly aligned temporal responses across individuals when they share similar interpretations of narratives (Leong et al., 2020; Yeshurun et al., 2017, 2021). In this context, the reduced neural synchrony observed in default mode network regions among IED-IED dyads suggests that individuals with IED may construct more idiosyncratic interpretations of the same social situations. Notably, similar patterns of idiosyncratic default mode network responses have been reported in lonely individuals (Baek et al., 2023), raising the possibility that individualized interpretations of social cues may be one contributing factor to maladaptive social relationships. The idiosyncratic interpretations of social contexts may also account for the observed effects in the dlPFC, a region known to support cognitive control, planning, and inhibition (Vanderhasselt et al., 2009). Prior work has shown that patients with IED exhibit heightened dlPFC activity during task-related error processing, suggesting impaired inhibitory regulation (Moeller et al., 2014). Accordingly, differences in how IED patients interpret social situations may give rise to differences in dlPFC engagement in the inhibition of aggression.

We also observed idiosyncratic responses in the insula and dorsal anterior cingulate cortex, two core regions of the salience network. The salience network is thought to support the integration of interoceptive signals and the detection of salient events, including potential threats, and to coordinate subsequent regulation of attention, emotion, and behavior (Barrett & Simmons, 2015; Craig, 2009; Fullana et al., 2016; Menon & Uddin, 2010; Uddin et al., 2017). Prior work has also found reduced dACC activation in individuals with high trait aggression during task-related frustration (Pawliczek et al., 2013). Reduced synchrony in these regions among IED participants may therefore reflect idiosyncratic threat perception and downstream regulatory processes that vary across individuals, which could contribute to aggressive behavior. Consistent with this interpretation, prior work has linked insula dysfunction to abnormalities in response control, socio-emotional information processing and elevated aggressive impulses (Blair, 2024; Coccaro et al., 2011; O’Nions et al., 2017). In this light, insula-related disruptions may reflect impaired aggression regulation in IED, serving as an affective mechanism through which idiosyncratic social interpretations translate into heightened reactive aggression.

By leveraging naturalistic video stimuli, the present study captured neural responses to dynamic, context rich social interactions. Our results suggest that IED does not reflect a single homogeneous neural profile, but instead is characterized by idiosyncrasy in how individuals process the same interpersonal cues. These findings align with recent work showing that individuals with maladaptive traits differ not only from healthy controls but also from others who share the same trait profile (Baek et al., 2023; Berman et al., 2013; Byrge et al., 2015; Liu et al., 2026; Yanagisawa et al., 2025), raising the possibility that neural idiosyncrasy reflects a transdiagnostic mechanism contributing to maladaptive social functioning. More broadly, naturalistic video paradigms combined with ISC-based measures may provide a useful framework for tracking changes in social information processing in response to treatment, as well as identifying neural targets for intervention aimed at improving socio-emotional regulation in real world interpersonal contexts.

The reduced neural synchrony in IED persisted after controlling for dyadic similarity in self-reported emotional responses and hostile intention attributions, suggesting that our results were not driven by idiosyncrasies in overt affective reactions or explicit judgments about others’ intent in the IED group. Instead, our findings point to variability in how social cues are integrated and interpreted, such that individuals with IED may arrive at distinct situation models even when they report similar emotions and comparable ratings of hostility. This interpretation-focused account may also explain why we did not observe effects in the amygdala, despite prior univariate evidence indicating heightened amygdala reactivity in IED (Coccaro et al., 2007), as amygdala activation may be more tightly coupled to the overall intensity of the affective response rather than interpretive processes. One limitation of this account is that the behavioral covariates used to control for participants’ intentionality attributions and emotional responses were based on self report ratings, which may not fully reflect the moment-to-moment interpretive processes that unfold during video viewing and contribute to neural synchrony. In addition, the study did not directly measure participants’ narrative interpretations of the social situations, which limits our ability to characterize the specific content of the idiosyncratic interpretations inferred from neural data. Future work could address these limitations by incorporating think aloud paradigms (Ke et al., 2025) and by applying content analytic approaches (Lyu et al., 2025) to more directly link the content of participants’ interpretations to patterns of neural idiosyncrasy during naturalistic social processing.

The present study demonstrates that IED individuals exhibit idiosyncratic neural responses during naturalistic social emotional processing. These findings suggest that IED is characterized not only by altered mean neural responses, but also by heightened variability in how individuals respond to the same social situations. By highlighting neural idiosyncrasy as a defining feature of social information processing in IED, this study highlights potential targets for interventions aimed at promoting healthier social cognition and reducing aggression in IED.

## Declarations

### Ethics approval

This study was approved by the IRB at the University of Chicago.

### Consent to participate

All subjects gave written informed consent.

### Consent for publication

All participants included in the study consented to the publication of the results in an anonymized form.

### Open Practices Statement

#### Availability of data and materials

The data are not publicly available as participants did not consent to releasing their data. The experiment was not preregistered.

#### Code availability

The analysis code for this study is publicly available on GitHub at https://github.com/ViCctorJJ/IED_Idiosyncrasy.

### Authors’ contributions

JC, SK, EFC and YCL conceived the study. SK and EFC collected the data. JC analyzed the data. JC and YCL interpreted the results. JC drafted the manuscript. JC, SK, EFC and YCL revised the manuscript. All authors read and approved the final manuscript.

