## Supplement materials for "Individuals with Intermittent Explosive Disorder Exhibit Idiosyncratic Neural Responses during Social-emotional Processing"

### Supplemental Methods

#### fMRIPrep Preprocessing Pipeline

Results included in this manuscript come from preprocessing performed using *fMRIPrep* 24.0.1 (Esteban et al., 2018, 2019; RRID:SCR_016216), which is based on *Nipype* 1.8.6 (Gorgolewski et al., 2011; Tustison et al., 2010; RRID:SCR_002502).

##### *Anatomical data preprocessing*

A total of 1 T1-weighted (T1w) images were found within the input BIDS dataset. The T1w image was corrected for intensity non-uniformity (INU) with N4BiasFieldCorrection (Tustison et al., 2010), distributed with ANTs 2.5.1 (Avants et al., 2008; RRID:SCR_004757), and used as T1w-reference throughout the workflow. The T1w-reference was then skull-stripped with a Nipype implementation of the antsBrainExtraction.sh workflow (from ANTs), using OASIS30ANTs as target template. Brain tissue segmentation of cerebrospinal fluid (CSF), white-matter (WM) and gray-matter (GM) was performed on the brain-extracted T1w using fast (FSL (version unknown), RRID:SCR_002823, Zhang et al., 2001). Volume-based spatial normalization to two standard spaces (MNI152NLin6Asym, MNI152NLin2009cAsym) was performed through nonlinear registration with antsRegistration (ANTs 2.5.1), using brain-extracted versions of both T1w reference and the T1w template. The following templates were were selected for spatial normalization and accessed with TemplateFlow (24.2.0, Ciric et al., 2022): FSL's MNI ICBM 152 non-linear 6th Generation Asymmetric Average Brain Stereotaxic Registration Model [Evans et al., 2012, RRID:SCR_002823; TemplateFlow ID: MNI152NLin6Asym], ICBM 152 Nonlinear Asymmetrical template version 2009c [10, RRID:SCR_008796; TemplateFlow ID: MNI152NLin2009cAsym].

***Functional data preprocessing***

For each of the 4 BOLD runs found per subject (across all tasks and sessions), the following preprocessing was performed. First, a reference volume was generated, using a custom methodology of fMRIPrep, for use in head motion correction. Head-motion parameters with respect to the BOLD reference (transformation matrices, and six corresponding rotation and translation parameters) are estimated before any spatiotemporal filtering using mcflirt (FSL, Jenkinson et al., 2002). The BOLD reference was then co-registered to the T1w reference using mri_coreg (FreeSurfer) followed by flirt (FSL, Jenkinson et al., 2002) with the boundary-based registration (Greve & Fischl, 2009) cost-function. Co-registration was configured with six degrees of freedom.

Several confounding time-series were calculated based on the preprocessed BOLD: framewise displacement (FD), DVARS and three region-wise global signals. FD was computed using two formulations following Power (absolute sum of relative motions, Greve & Fischl, 2009) and Jenkinson (relative root mean square displacement between affines, Jenkinson et al., 2002). FD and DVARS are calculated for each functional run, both using their implementations in Nipype (following the definitions by Power et al. (2014)). The three global signals are extracted within the CSF, the WM, and the whole-brain masks. Additionally, a set of physiological regressors were extracted to allow for component-based noise correction (CompCor, Behzadi et al., 2007). Principal components are estimated after high-pass filtering the preprocessed BOLD time-series (using a discrete cosine filter with 128s cut-off) for the two CompCor variants: temporal (tCompCor) and anatomical (aCompCor). tCompCor components are then calculated from the top 2% variable voxels within the brain mask. For aCompCor, three probabilistic masks (CSF, WM and combined CSF+WM) are generated in anatomical space. The implementation differs from that of Behzadi et al. in that instead of eroding the masks by 2 pixels on BOLD space, a mask of pixels that likely contain a volume fraction of GM is subtracted from the aCompCor masks. This mask is obtained by thresholding the corresponding partial volume map at 0.05, and it ensures components are not extracted from voxels containing a minimal fraction of GM.

Finally, these masks are resampled into BOLD space and binarized by thresholding at 0.99 (as in the original implementation). Components are also calculated separately within the WM and CSF masks. For each CompCor decomposition, the k components with the largest singular values are retained, such that the retained components' time series are sufficient to explain 50 percent of variance across the nuisance mask (CSF, WM, combined, or temporal). The remaining components are dropped from consideration. The head-motion estimates calculated in the correction step were also placed within the corresponding confounds file. The confound time series derived from head motion estimates and global signals were expanded with the inclusion of temporal derivatives and quadratic terms for each (Satterthwaite et al., 2013). Frames that exceeded a threshold of 0.5 mm FD or 1.5 standardized DVARS were annotated as motion outliers. Additional nuisance timeseries are calculated by means of principal components analysis of the signal found within a thin band (crown) of voxels around the edge of the brain, as proposed by Patriat et al. (2017). All resamplings can be performed with a single interpolation step by composing all the pertinent transformations (i.e. head-motion transform matrices, susceptibility distortion correction when available, and co-registrations to anatomical and output spaces). Gridded (volumetric) resamplings were performed using nitransforms, configured with cubic B-spline interpolation.

Many internal operations of fMRIPrep use Nilearn 0.10.4 (Abraham et al., 2014, RRID:SCR_001362), mostly within the functional processing workflow. For more details of the pipeline, see the section corresponding to workflows in fMRIPrep's documentation.

***Copyright Waiver***

The above boilerplate text was automatically generated by fMRIPrep with the express intention that users should copy and paste this text into their manuscripts unchanged. It is released under the CC0 license.

**Bayesian model specification**

We fit a Bayesian multilevel model to predict dyadic ISC values (Fisher z-transformed correlation *cor_fz*). The primary model included dyadic diagnostic composition as the main predictor, together with two dyadic control variables indexing behavioral similarity in attribution and emotional response. To account for the non-independence of dyadic observations, we included a multi-membership random intercept for the two members of each dyad. In brms syntax, the primary model was specified as:

*cor_fz* = 1 + *x_ij* + ATTR_isc + ER_isc + (1 | mm(sub1, sub2))

Here, *x_ij* denotes dyadic diagnostic composition (i.e., dyad type), which was entered as a categorical predictor. ATTR_isc and ER_isc represent dyadic similarity in attribution and emotional response, respectively. The term (1 | mm(sub1,sub2)) specifies a multi-membership random intercept to account for the fact that each dyadic observation is jointly associated with two participants.

In an additional covariate-adjusted model, we further controlled for the race of each dyad member by including two subject-level race covariates as fixed effects:

*cor_fz* = 1 + *x_ij* + ATTR_isc + ER_isc + race1 + race2 + (1 | mm(sub1, sub2)) where race1 and race2 denote the race category of the first and second member of the dyad, respectively.

***Prior specification***

For population-level effects, including the categorical effect of dyadic diagnostic composition and the covariates ER_isc, ATTR_isc, and, where applicable, race1 and race2, we used the default improper flat priors for population-level effects (Chen et al., 2020).

For the standard deviation of the subject-level multi-membership random intercept, we specified a weak Student-*t* prior (Chen et al., 2020):

$$\mathrm{SD}_{\mathrm{subject}} \sim \text{Student-}t(3,0,1)$$

Because standard deviation parameters are constrained to be positive, this corresponds to a half-Student-*t* prior.

For the residual standard deviation, we specified a Cauchy prior centered at zero with scale equal to the empirical standard deviation of *cor_fz*:

$$\sigma \sim \mathrm{Cauchy}(0, sd_{cor\_fz})$$

Because $\sigma > 0$, this corresponds to a half-Cauchy prior on the residual standard deviation.

#### Supplemental Results

| Current syndromal disorders | n |
|---|---|
| Any depressive mood disorder | 5 (26.3%) |
| Any anxiety disorder | 4 (21.2%) |
| Traumatic and stress disorders | 2 (10.5%) |
| Impulse control disorders (Not-IED) | 1 (5.3%) |
| **Lifetime syndromal disorders** | |
| Any depressive mood disorder | 10 (52.6%) |
| Substance use disorder | 9 (47.4%) |
| Any anxiety disorder | 6 (31.6%) |
| Traumatic and stress disorders | 4 (21.1%) |
| Eating disorder | 1 (5.3%) |
| Impulse control disorders (Not-IED) | 2 (7.7%) |
| **Personality disorders** | |
| Any personality disorder | 16 (84.2%) |
| Cluster A (Odd) | 4 (21.2%) |
| Cluster B (Dramatic) | 3 (15.8%) |
| Cluster C (Anxious) | 1 (5.3%) |
| PD-NOS | 11 (57.9%) |

**Table S1.** DSM-5 syndromal and personality disorders in the IED group (Coccaro et al., 2022).

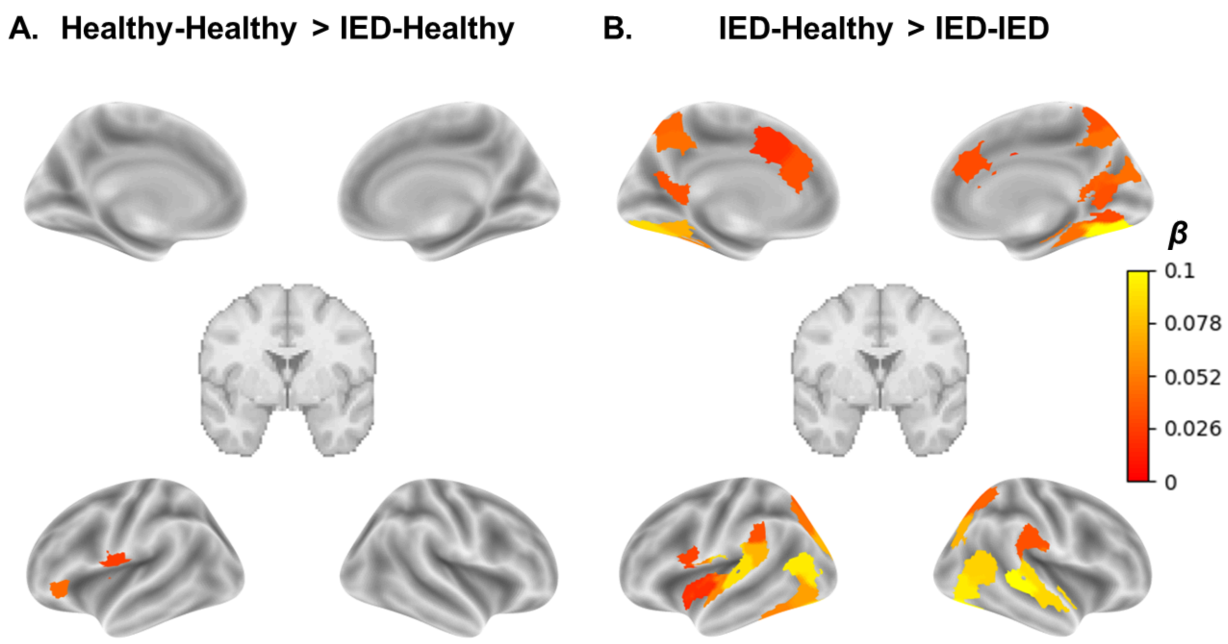


**Figure S1** ROIs showing significant differences in dyadic ISC across groups (color bar indicates the group difference ($\beta$), i.e. the estimated posterior means of dyad type). The two subplots use the same color scale. The models controlled for dyadic similarity in attribution and emotional responses.

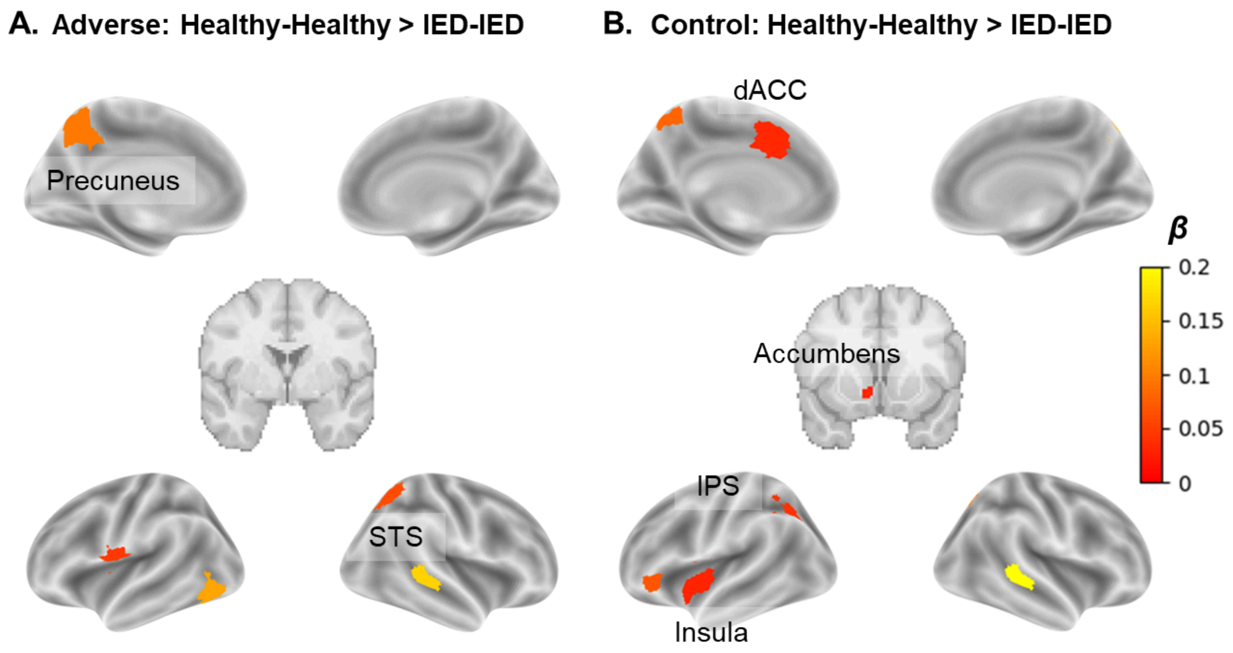


**Figure S2** ROIs showing significant differences in dyadic ISC between HH and II groups (color bar indicates the group difference ($\beta$), i.e. the estimated posterior means of dyad type) when participants were watching adverse or control videos. The two subplots use the same color scale. The models controlled for dyadic similarity in attribution and emotional responses.

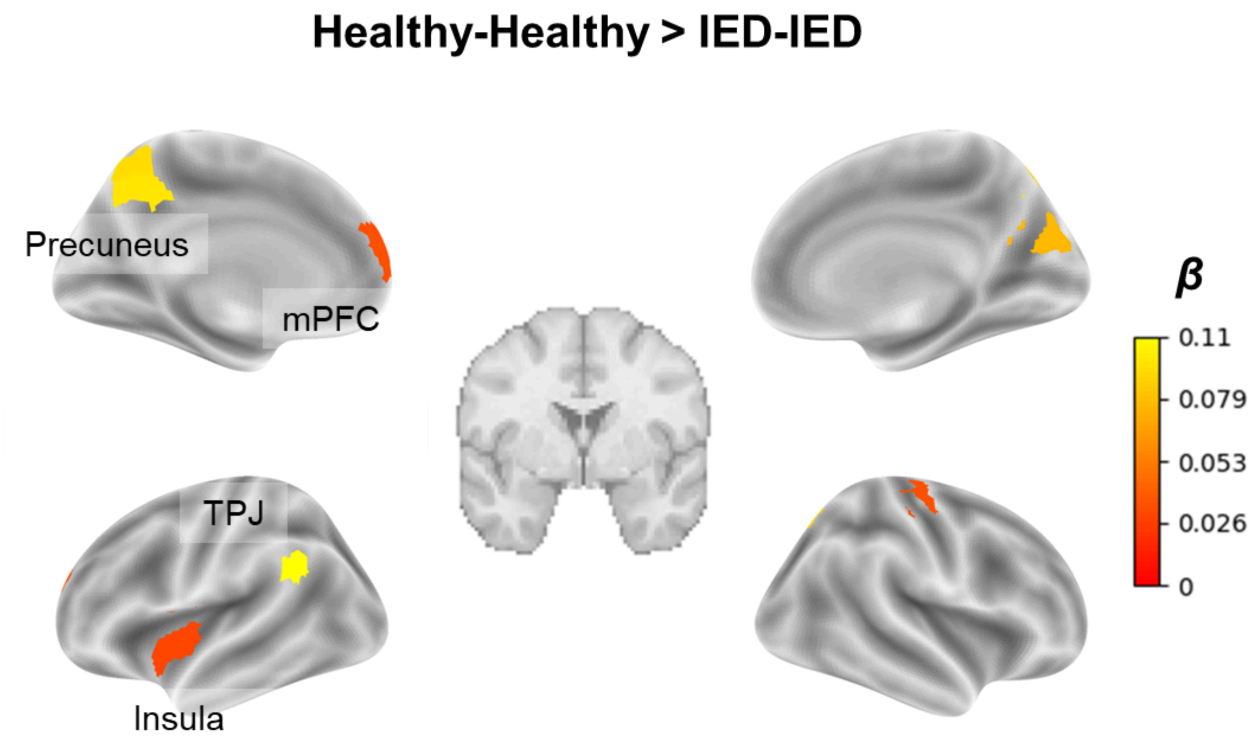


**Figure S3** ROIs showing significant differences in dyadic ISC between HH and II groups (color bar indicates the group difference ($\beta$), i.e. the estimated posterior means of dyad type) during video watching while controlling for the racial composition of the dyad.. The two subplots use the same color scale. The models also controlled for dyadic similarity in attribution and emotional responses.
